# Cell-type specific open chromatin profiling in human postmortem brain infers functional roles for non-coding schizophrenia loci

**DOI:** 10.1101/062513

**Authors:** John F. Fullard, Claudia Giambartolomei, Mads E. Hauberg, Ke Xu, Christopher Bare, Joel T. Dudley, Manuel Mattheisen, Joseph K. Pickrell, Vahram Haroutunian, Panos Roussos

**Author notes:** **Correspondence:** Dr. Panos Roussos, Icahn School of Medicine at Mount Sinai, Department of Psychiatry and Department of Genetics and Genomic Science and Institute for Multiscale Biology, One Gustave L. Levy Place, New York, NY, 10029, USA.

## Abstract

To better understand the role of *cis* regulatory elements in neuropsychiatric disorders we applied ATAC-seq to neuronal and non-neuronal nuclei isolated from frozen postmortem human brain. Most of the identified open chromatin regions (OCRs) are differentially accessible between neurons and non-neurons, and show enrichment with known cell type markers, promoters and enhancers. Relative to those of non-neurons, neuronal OCRs are more evolutionarily conserved and are enriched in distal regulatory elements. Our data reveals sex differences in chromatin accessibility and identifies novel OCRs that escape X chromosome inactivation, with implications for intellectual disability. Transcription factor footprinting analysis identifies differences in the regulome between neuronal and non-neuronal cells and ascribes putative functional roles to 16 non-coding schizophrenia risk variants. These results represent the first analysis of cell-type-specific OCRs and TF binding sites in postmortem human brain and further our understanding of the regulome and the impact of neuropsychiatric disease-associated genetic risk variants.

## INTRODUTION

Epigenetic modification of chromatin, including its 3-dimensional structure, plays a central role in the regulation of gene expression and is essential for development and the maintenance of cell identity and function (Fullard et al.). A large proportion of neuropsychiatric disease-associated loci, such as those in schizophrenia (SCZ) (PGC-SCZ, 2014) and Alzheimer’s disease (AD) (Lambert et al., 2013), are non-coding and, as such, have no impact on the structure of proteins. Instead, these non-coding risk variants are thought to exert their effects by altering the function of *cis* regulatory elements (CREs) required for the correct spatiotemporal expression of genes (Maurano et al., 2012; Roadmap Epigenomics et al., 2015; Roussos et al., 2014; Trynka et al., 2013). Importantly, *cis* regulation of gene expression is often specific for tissue and even cell type (Maurano et al., 2012; Roadmap Epigenomics et al., 2015). Correspondingly, disease-associated loci have been shown to be enriched for CREs in tissues and cells relevant to the pathophysiology of disease (Maurano et al., 2012; Roussos et al., 2014; Trynka et al., 2013).

Employing existing epigenome data to further our understanding of neuropsychiatric diseases is hindered by the fact that much of the epigenomic landscape remains unexplored in the relevant cells: The Encyclopedia of DNA Elements (ENCODE) consortium (Bernstein et al., 2012; Maurano et al., 2012), Roadmap Epigenomics Mapping Consortium (REMC) (Roadmap Epigenomics et al., 2015; Zhu et al., 2013), and FANTOM5 (Andersson et al., 2014) all focused on actively dividing cells or used only homogenate brain tissue. A limitation to the latter approach is that the resultant data was derived from a mixture of markedly different cells such as neurons, microglia, oligodendrocytes, and astrocytes. Because CRE-mediated epigenetic regulation shows cell-type specificity (Maurano et al., 2012; Roadmap Epigenomics et al., 2015), the study of mixed cell populations can fail to detect cell-type-specific signals. Furthermore, in studies using homogenized brain tissue, the proportion of each cell type contributing to the assay is undetermined, further increasing sample-to-sample variability.

Here we present, to our knowledge, the first cell-type-specific map of OCRs in human frontopolar prefrontal cortex. We used the Assay for Transposase Accessible Chromatin followed by sequencing (ATAC-seq) to map open chromatin regions (OCRs) (Buenrostro et al., 2013). We generated and analyzed 3.1 billion reads from 2 broad cell types (neuronal and non-neuronal) isolated from frozen postmortem tissue of persons with no known neuro-or psycho-pathology by fluorescence-activated nuclear sorting (FANS) in 8 samples, and identified thousands of cell-type specific OCRs. We further identified several OCRs that escape X chromosome inactivation in a cell-type specific manner and
are implicated in X-linked intellectual disability. Additionally, we inferred transcription factor (TF) binding using footprinting to interrogate cell-type differences in the regulation of gene expression. Finally, a number of the OCRs and TF binding sites identified by ATAC-seq coincide with known SCZ risk loci, allowing us to pin-point the likely causative SNP therein and providing evidence for a putative mechanism by which these variants contribute to the etiology of SCZ.

## RESULTS

### Profiling chromatin accessibility in Neuronal and non-Neuronal Cells

We assessed the landscape of OCRs in neuronal and non-neuronal nuclei derived from frozen postmortem frontopolar prefrontal cortex [Brodmann area 10 (BA10)] of eight samples (**Figure 1A**). We chose BA10 as it is a brain region that is important for cognition (Gilbert et al., 2006) and has been associated with neuropsychiatric diseases (Takizawa et al.). For each sample, ATAC-seq was performed on 50,000 neuronal (NeuN positive) and non-neuronal (NeuN negative) nuclei isolated by FANS (**Supplemental Experimental Procedures**). To control for potential sample confounds, we used samples from a single brain bank isolated from individuals with similar genetic background (all Caucasians) that had not been diagnosed with neurological or psychiatric conditions at the time of death (**Table 1**) and evidenced no discernable neuropathology upon detailed postmortem examination (Purohit et al., 1998). To control for potential experimental factors, all steps were performed at one site and samples were randomized prior to FANS and sequencing. We assessed the effects of technical variability on ATAC-seq profiles by comparing the results obtained from triplicates derived from one neuronal and one non-neuronal sample isolated from BA10. We found a high concordance of read coverage (number of reads) among replicates, within genomic bins of 1000bp (average Pearson Correlation Coefficient *R* (PCC) = 0.97) (**Figure S1A and A’**).

**Table 1.** Clinical characteristics of the postmortem cohort used in this study.

| Sample | Gender | Age of death<br>(years) | Postmortem<br>interval (hours) | Brain weight<br>(grams) | Tobacco<br>use | Cause of death | Hypertension | Diabetes (Insulin<br>dependent) | Diabetes (Non<br>insulin dependent) |
| --- | --- | --- | --- | --- | --- | --- | --- | --- | --- |
| S1 | Female | 85 | 5 | 1128 | No | Cardio respiratory failure | No | No | No |
| S2 | Male | 64 | 23.3 | 1238 | Yes | Cardio respiratory failure | Yes | No | No |
| S3 | Male | 79 | 13 | 1130 | Yes | Lung cancer | No | No | No |
| S4 | Male | 75 | 5 | 1276 | No | Myocardial infarction | Yes | No | Yes |
| S5 | Female | 90 | 5.9 | 984 | No | Cardio respiratory failure | Yes | No | Yes |
| S6 | Female | 90 | 1.8 | 876 | No | Atrial fibrillation | Yes | No | No |
| S7 | Female | 85 | 8 | 1040 | No | Cardio respiratory failure | Yes | No | No |
| S8 | Male | 93 | 3.5 | 1104 | No | Cardio respiratory failure | Yes | No | No |

The average number of uniquely mapped and non-duplicated paired-end reads per sample was 54.8 million, with low mitochondrial DNA contamination (average 0.7% of the mapped reads) (**Table S1**). There was no significant difference in the number of reads among neuronal (50.7 million reads) and non-neuronal (58.8 million reads) samples (Student’s t-test P = 0.5). We applied the irreproducible discovery rate (IDR) approach in the technical replicates and chose MACS2 as the best peak-calling algorithm (**Figure S1B**) with P = 4.7 × 10^−6^ as the MACS2 cutoff for calling peaks at IDR 5% (**Figure S1B’**). Using these parameters, the average number of called peaks per sample was 47,911, with no difference between neuronal (50,483 peaks) and non-neuronal (45,338 peaks) samples (Student’s t-test P = 0.6). (**Table S1**). A total of 72,033 and 68,606 peaks were defined when the filtered peaks called from neuronal and non-neuronal samples, respectively, were combined (**Figure S1C and C’**). Approximately one third of neuronal and non-neuronal peaks overlapped (**Figure 1B**).

**Figure 1.**
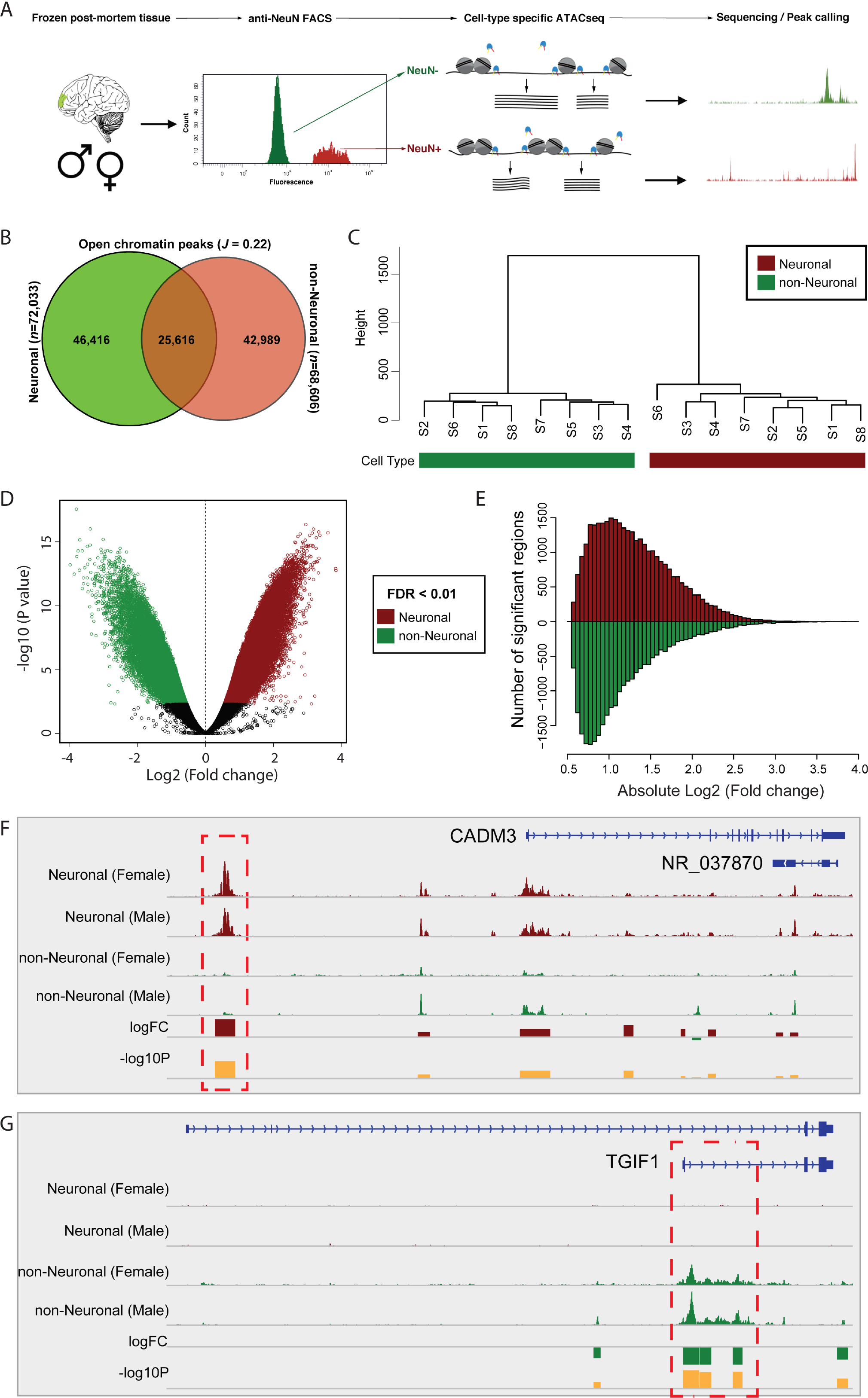
Differential analysis of chromatin accessibility in neuronal and non-neuronal cells. **(A)** Schematic outline of study design. (**B)** Venn diagram showing the overlap of neuronal and nonneuronal OCRs. **(C)** Unsupervised hierarchical clustering of ATAC-seq data. The normalized read count matrix across 151,021 peak regions is shown. **(D)** Volcano plot showing the distribution of −log10 p-value and log2 fold-change of differential chromatin accessibility analysis across 151,021 OCRs. **(E)** Distribution of log2 fold-change of differential chromatin accessibility analysis for 33,054 neuronal and 27,599 non-neuronal OCRs. Averaged cell-type and gender-specific ATAC-seq signals at **(F)** *CADM3* and **(G)** *TGIF1* gene loci. Positive and negative logFC indicate neuronal and nonneuronal differential signals, respectively. Red box indicates the significant differentially accessible region in **(F)** neuronal and **(G)** non-neuronal cells.

For quantitative analysis of differences among neuronal and non-neuronal samples, we generated a matrix of mapped reads using a final consensus of 115,021 peaks (**Supplemental Experimental Procedures**). Unsupervised hierarchical clustering of the normalized mapped reads in each peak region identified a clear distinction between neuronal and non-neuronal samples. (**Figure 1C**). Exploratory analysis identified confounds (age at death, gender and postmortem interval) as significant predictors of chromatin accessibility (**Figure S1D and E**). Following data normalization of read counts for each peak, a quasi-likelihood negative binomial generalized model, adjusting for confounds, was performed (**Supplemental Experimental Procedures**). We identified 60,653 differentially modified OCRs (adjusted false discovery rate (FDR) < 0.01, **Figure 1D**, **Table S2**). Among these, 33,054 were neuronal and 27,599 were non-neuronal, with a moderate to large average fold change (FC) (median FC 2.19; range 1.46-15.86, **Figure 1E**). For example, a neuron-specific regulatory region (chr1:159110612-159112536; log_2_FC = 2.89, P = 2.76 × 10^−23^, adjusted P = 9.1 × 10^−11^) is positioned ~29kb upstream of CADM3 (Cell adhesion molecule 3), a gene that is highly expressed in cortical pyramidal cells (Zeisel et al., 2015) (**Figure 1F**). A non-neuronal region (chr18:3447649-3448767; log2FC = −3.99, P = 6.46 × 10^−14^, adjusted P = 4.13 × 10^−11^) is positioned within the transcription start site (TSS) of the shorter TGIF1 (TGFB-Induced Factor Homeobox 1) isoform, a gene with high expression in microglia (Zhang et al., 2014) associated with Holoprosencephaly-4 (OMIM: 142946) (**Figure 1G**).

### Annotation of the Cell-type Specific Regulome in Neuronal and non-Neuronal Cells

We analyzed the distribution of all and differentially accessible elements (neuronal and non-neuronal) as a function of the distance from the nearest TSS. For all three categories (all, neuronal and nonneuronal) we observed an enrichment of OCRs proximal to TSSs (**Figure 2A**). Relative to all OCRs, we observed a depletion of neuronal OCRs (OR = 0.7, P = 9.8×10^−39^) and an enrichment of nonneuronal OCRs (OR = 1.3, P = 1.2×10^−37^) in the vicinity of TSSs (**Figure 2B**). There was, however, enrichment for neuronal OCRs located 10kb – 100kb downstream of TSSs (OR = 1.3, P = 1.3×10^−127^). Next, we examined the enrichment of all and differentially accessible elements in terms of genomic features. In agreement with our findings that neuronal accessible regions are relatively depleted in the proximity of TSSs, we identified a depletion for neuronal OCRs located in promoters (OR =0.7, P = 8.2 × 10^−113^) and an enrichment in intronic regions (OR =1.2, P = 6.5 × 10^−130^) (**Figure 2C**), when compared to non-neurons. Overall, relative to non-neuronal OCRs, the differentially accessible regions of neurons are enriched for distal regulatory elements, suggesting a more important role for long-range regulation of gene expression in neurons.

We next examined if the identified OCR regions were under evolutionary constraint as evidenced by higher Genomic Evolutionary Rate Profiling (GERP) scores (Cooper et al., 2005). When averaging across the OCRs, we found a peak in the GERP score coinciding with the center of the OCR for both the neuronal and non-neuronal cells (**Figure 2D**). Interestingly, neuronal OCRs exhibit markedly higher GERP scores than non-neuronal OCRs (0.26 vs 0.14, p = 7.7 × 10^−14^ in a two-sided t test). To further explore this observation, and to correct for potential confounders, we stratified the analysis based on OCRs found in exons, promoters, introns, and intergenic regions (**Figure S2**). Compared to non-neuronal OCRs, neurons show higher GERP score in intergenic and promoter regions (two-sided t test p < 2.2 × 10 for both comparisons), respectively. Therefore, it appears that neuronal OCRs in distal and proximal regulatory regions are under stronger functional constraint than those of other brain cells.

We compared the ATAC-seq peaks with published histone modification data from homogenates of human prefrontal cortex (Roadmap Epigenomics et al., 2015) (**Figure 2E**). The ATAC-seq peaks (all peaks or cell type-specific peaks) showed enrichment for active and poised promoters and active and repressed enhancers. No enrichment for transcribed, polycomb-repressed or heterochromatin regions was observed. Systematic comparison of cell-type specific OCRs with published regulatory sequences for enhancers and promoters across multiple tissues showed a strong enrichment in neuronal OCRs for brain-related enhancers and promoters (**Figure S3A and S3B**). Convincingly, the strongest enrichment across all enhancers for neuronal OCRs was observed for prefrontal cortex. The non-neuronal OCRs show lower specificity for brain related enhancers and promoters. Finally, a comparison with DNase I hypersensitivity sequencing (DHS-seq) regions across 39 tissues show that brain is the most enriched tissue in our data set (**Figure S3C**). Overall, the analysis shows our cell-type-specific ATAC-seq data is enriched with previously annotated regulatory sequences for promoters, enhancers and OCRs in human brain tissue.

**Figure 2.**
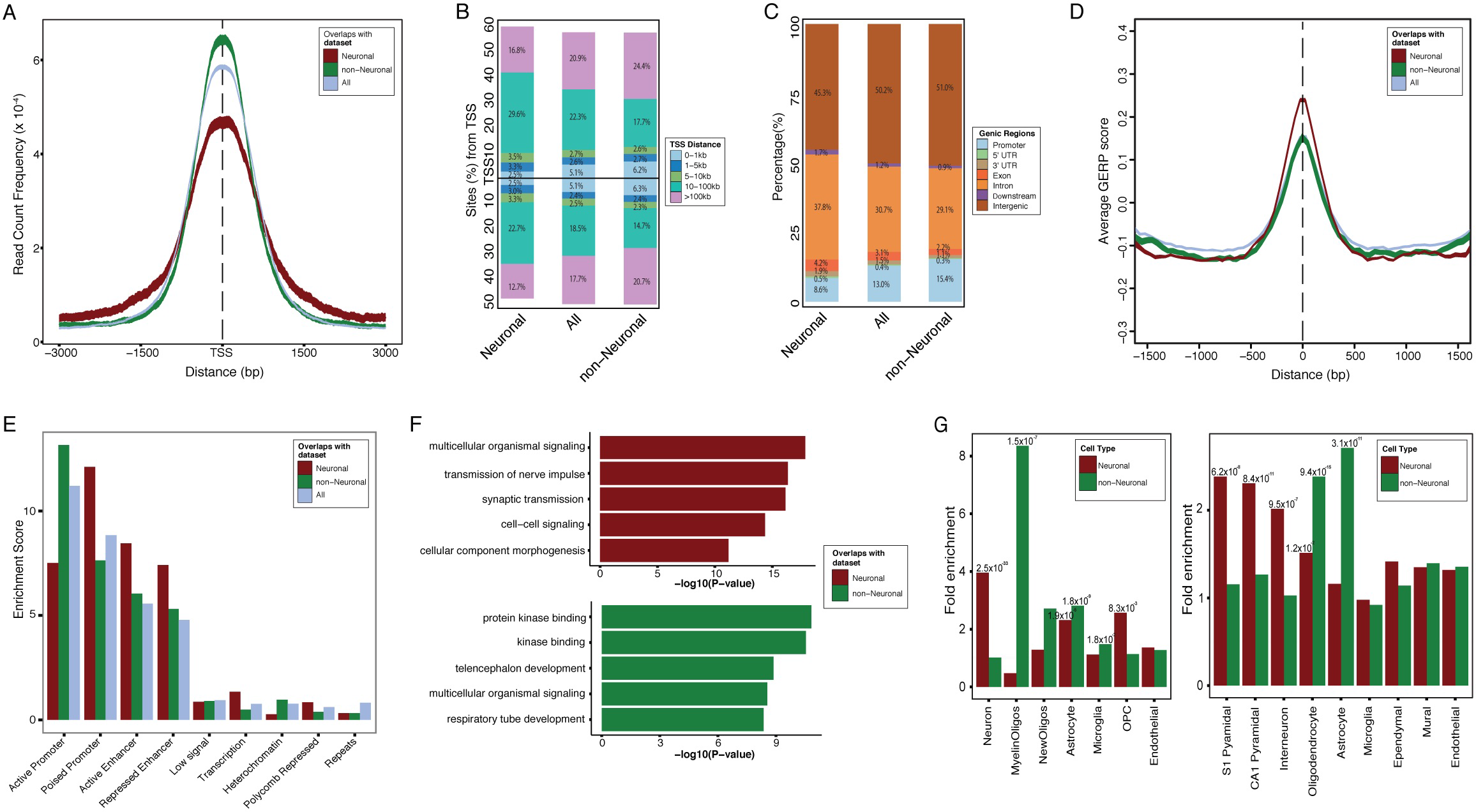
Annotation of the neuronal and non-neuronal regulome. **(A)** Average read count frequency of OCRs in TSS regions. Confidence interval estimated by bootstrap method. **(B)** Distribution of all peaks and differential OCRs relative to TSS. **(C)** Distribution of genomic features of all and differential OCRs. **(D)** Average GERP score as a function of distance from the center of [−1000bp, 1000bp] of all peaks and differential OCRs. Curves and their 95% confidence intervals are calculated on a 50 bp sliding window. (**E)** Enrichment of all peaks and differential OCRs in various human prefrontal brain tissue chromatin states. **(F)** Gene Ontology terms for differential OCRs enriched in neuronal (dark red) and non-neuronal (dark green) samples. **(G)** Enrichment of differential OCRs for neuronal (dark red) and non-neuronal (dark green) cell type-specific markers.

To further explore the identified OCRs, we conducted pathway analysis by annotating peaks to proximal genes. In this analysis, neuronal peaks showed a significant enrichment for the terms synaptic transmission, cell-cell signaling, and cellular morphogenesis (**Figure 2F**), whereas nonneuronal peaks were enriched for pathways related to protein kinase binding and telencephalon development. Next, we examined the overlap between genes proximal to the differentially accessible elements and cell type-specific markers derived from mouse brain studies (Zeisel et al., 2015; Zhang et al., 2014). We detected a highly significant overlap between the set of the predicted human neuronal genes and the set of cell type-specific genes for neurons (Zhang et al., 2014) (P = 2.5 × 10^33^) or pyramidal cells (S1 pyramidal markers: P = 6.2 × 10^−8^; CA1 pyramidal markers: P = 8.4 × 10^−11^) and interneurons(Zeisel et al., 2015) (P = 9.5 × 10^−7^) in the mouse brain (**Figure 2G**). For nonneuronal OCRs, we observed a strong enrichment for oligodendrocytes (P < 1.5 × 10^−7^) and astrocytes (P < 1.8 × 10^−9^) in both studies (Zeisel et al., 2015; Zhang et al., 2014).

### Sex-Specific chromatin accessibility in Neuronal and non-Neuronal Cells

Although females have two X chromosomes and males only one, most of the genes on this chromosome show similar levels of expression across sexes. This widespread dosage compensation results from a phenomenon called X chromosome inactivation (XCI), which involves XIST and other long non-coding RNAs (Lee and Bartolomei, 2013). However, a subset of X-linked genes escape XCI through a poorly understood mechanism, and for some genes this occurs in a tissue and cell type specific manner (Deng et al., 2014). Using our OCR data we were able to interrogate such XCI escapees along with other gender-specific variations in chromatin structure as described below.

Not surprisingly, the majority of differential OCRs between males and females mapped to the sex chromosomes **(Figure 3A and Table S3)**. For neuronal cells, 225 out of 343 gender specific OCRs (66%), at FDR 1%, mapped to the sex chromosomes compared to 291 out of 347 OCRs (84%) at FDR 1% in non-neuronal cells (Figure 3B). Only male subjects showed signal for neuronal and nonneuronal OCRs on the Y chromosome, accounting for all accessible regions for the Y chromosome **(Figure 3A and 3B)**. The majority of OCRs on the X chromosome are inactivated in females and only 3.9% (154 out of 3,947 regions) are more accessible compared to males **(Figure 3A and 3B)**, indicating the presence of OCRs on both copies of the X chromosome. Pathway analysis of female-and male-biased accessible regions showed significant enrichment for gender-specific biological
processes and molecular functions **(Figure 3C)**. Interestingly, female-biased OCRs showed enrichment for gene sets related to intellectual disability and autism.

**Figure 3.**
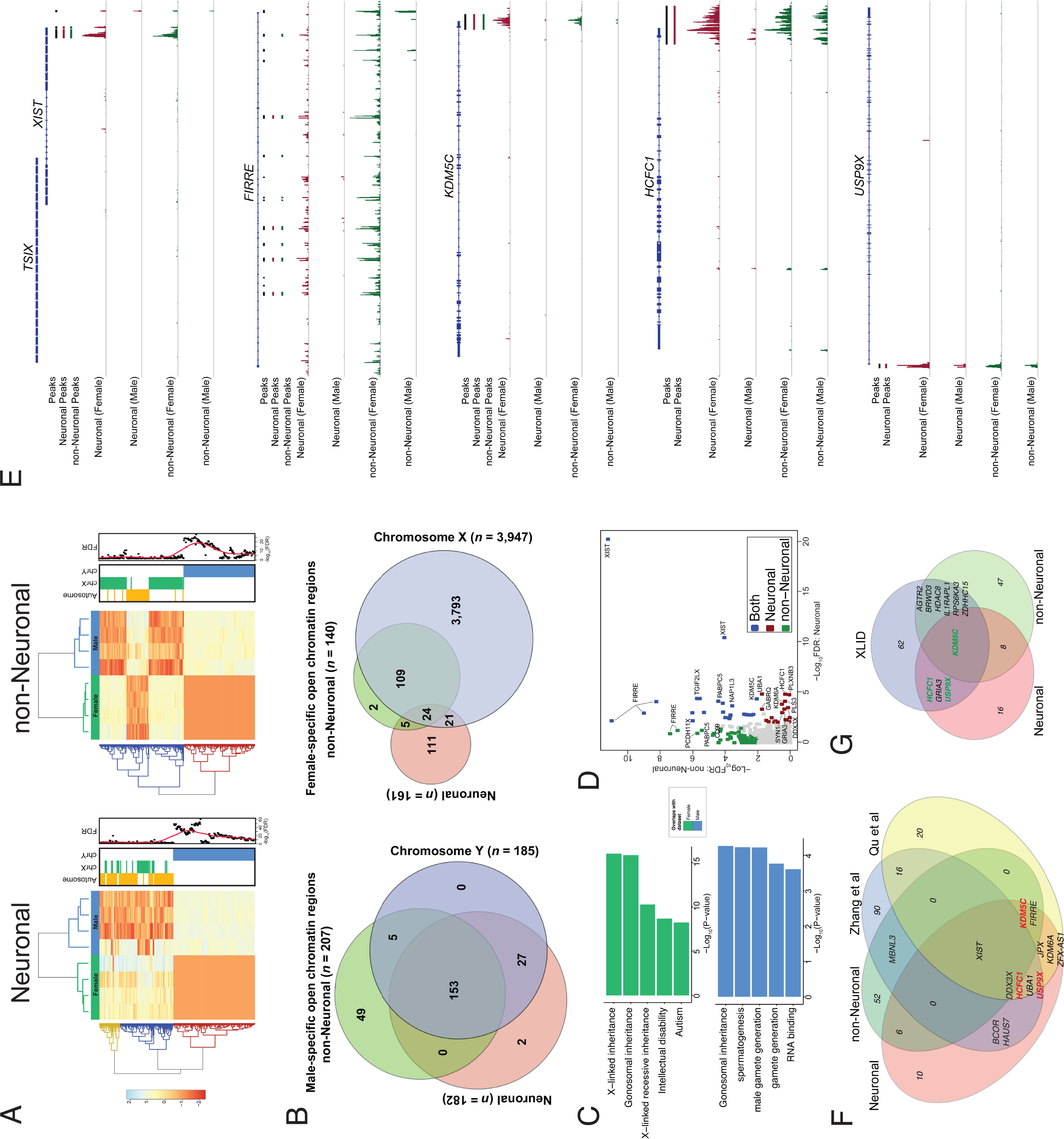
Gender-specific regulome in neuronal and non-neuronal cells. **(A)** Association of gender-specific regulatory activity in neuronal (left) and non-neuronal cells (right). Left: Bivariate clustering of samples (columns) and OCRs (rows) depicts the gender-specific regulatory activity in neuronal and non-neuronal samples, as marked by the dark green (female) and blue (male) horizontal bar at top. Color scale indicates relative ATAC-seq signal as indicated. Middle: bar graph indicates regulatory elements on autosomes (orange), chromosomes X (chrX; green) and Y (chrY; blue). Right: FDR of significance for each regulatory element. **(B)** Venn diagrams showing the distribution of gender specific differential OCRs in neuronal and non-neuronal samples for elements on chromosomes Y (left) and X (right). **(C)** Gene Ontology terms enriched in female (dark green) and male (blue) differential OCRs. **(D)** Scatterplot showing the distribution of gender-specificregulatory activity in neuronal and non-neuronal cells across 3,947 OCRs on chromosome X. The x-axis and y-axis indicate the −log10 FDR from the gender-specific differential analysis. Both (blue color) indicates OCRs that have FDR < 1% in both neuronal and non-neuronal samples; Neuronal (dark red) indicate significant OCRs only in neuronal cells and log2 fold chance > 1.25; non-Neuronal (forest green) indicate significant OCRs only in non-neuronal cells and log2 fold chance > 1.25. **(E)** Averaged cell-type and gender-specific ATAC-seq signals at *XIST*, *FIRRE*, *KDM5C*, *HCFC1* and *USP9X* gene loci. Peaks indicate the position of OCRs; neuronal and non-neuronal peaks indicate gender-specific differential accessible regulatory elements at FDR 1%. **(F)** Venn diagram showing the intersection of genes predicted to escape chromosome X inactivation (XCI) in neuronal and nonneuronal samples and known escapee genes from two previous studies (Zhang et al and Qu et al). Gene symbols are presented for genes that have been reported previously and those that have been associated with X-linked intellectual disability (XLID) are highlighted (red). **(G)** Venn diagram showing the intersection of genes predicted to escape XCI in neuronal and non-neuronal samples and genes that have been associated with XLID. Known escapee genes based on Zhang et al and Qu et al studies are highlighted (green).

We identified 45 neuronal and 133 non-neuronal OCRs associated, respectively, with 28 and 62 genes that escape XCI (**Figure 3D**). Of these 90 genes, only nine were common, indicating cell type-specific XCI in human brain cells. The most significant OCR in both neuronal and non-neuronal cells is in the promoter region of XIST, an X-encoded gene that, as previously mentioned, plays a central role in XCI and is only active in cells of females (**Figure 3D** and **3E**). Out of the 81 genes we predict to escape XCI in neuronal and non-neuronal cells, a subset of 13 genes – 9 neuronal, 1 non-neuronal and 3 in both – are genes known, or have been predicted previously, to escape XCI (Qu et al.; Zhang et al., 2013) (**Figure 3F**), including *FIRRE*, a nuclear-organization lncRNA that interacts with, and influences, higher-order nuclear architecture across chromosomes (Hacisuleyman et al., 2014; Yang et al., 2015). In agreement with previous ATAC-seq data in human CD4+ T cells (Qu et al.), we identified nine *FIRRE* enhancers in introns 2-12 which are active in female non-neuronal cells, while only 4 are active in female neuronal cells, indicating a larger number of non-neuronal OCRs that escape XCI relative to neuronal cells (**Figure 3E**). We then examined if any of the genes identified as escaping XCI have been implicated in X-linked intellectual disability (XLID category from HUGO Gene Nomenclature Committee). Of the 81 genes predicted to escape XCI in neuronal and non-neuronal cells, 10 have previously been associated with XLID (**Figure 3G**), consisting of 3 known escapee genes (*KDM5C*, *USP9X* and *HCFC1*) (**Figure 3E**) and 7 novel genes, including the glutamate ionotropic receptor AMPA type subunit 3 (*GRIA3*). Overall, these results reveal sex differences in chromatin accessibility and identify novel, cell-type specific, OCRs that escape XCI and are in proximity to XLID-associated genes.

### Cell-type Specific Transcription factor binding and gene regulation

As most transcription factors (TFs) bind preferentially to open chromatin (Thurman et al., 2012) it is of interest to analyze such regions with regard to TF binding. We did so with a set of 432 motifs representing 864 TFs aggregated from multiple databases (Weirauch et al., 2014) (**Supplemental Experimental Procedures**). Out of these motifs, 179 show a fold enrichment of binding sites ≥ 1.25 in the peaks of either the neuronal or the non-neuronal samples compared to the genomic background. We chose to primarily focus on these motifs. In addition to being enriched in motif binding sites, the shape of the reads within ATAC-peaks can also be used to infer TF binding, as illustrated for the TF CTCF (**Figure 4A**). Here, the binding sites are surrounded by a characteristic pattern of transposition insertion while the actual binding site shows a “footprint” as the bound TF sterically protects the DNA from transposase integration (Buenrostro et al., 2013; Neph et al., 2012).

To exploit the pattern of transposition insertion in order to improve TF binding prediction, we applied the protein interaction quantitation (PIQ) (Sherwood et al., 2014) framework, which, for each motif, learns the pattern of transposition insertion in the vicinity of potential binding sites. To measure the relative importance of each TF in the neuronal and non-neuronal samples, we used the output of PIQ to calculate motif scores (**Supplemental Experimental Procedures**) and compared them against each other (**Figure 4B**). We note that similar results to the one described below could be obtained by comparing the enrichment of motifs in the ATAC-seq peaks regardless of footprinting (**Figure S4**). To quantify the difference between the two cell types, we used the ratio of the motif scores to categorize the motifs as showing probable, or definite, cell-type specificity (**Figure 4B, Table S4, Table S5**).

The most prominent cell-type-specific TF differences was observed for promoter-depleted TFs. The TFs most specific for neurons were the FOS/JUN families (which together form the AP1-complex), showing a 2.7-fold higher motif score in neuronal cells. These TFs also showed greater than two-fold enrichment in neuronal peaks with, conversely, a slight depletion in peaks of non-neurons (**Table S5**). The RFX-family represented the next-most neuronal specific TF motif. These findings complement a recent study that identified these two groups of TFs as central regulators of excitatory neuronal function (Mo et al., 2015). For non-neuronal cells, the two most specific motifs were ONECUT1/2/3 and PAX3/7, of which the latter has the most established role in brain function (Baldwin et al., 1995; Seo et al., 1998). We further note that the TF motifs specific to either the neuronal or non-neuronal cells appear to fall into distinct families. For instance, the Basic helix-loop-helix (bHLH) family seems to be primarily active in neuronal cells whereas the homeodomain/sox TF families seem to be predominantly non-neuronal (**Table S4**). On the other hand, several motifs of general TFs are identified mostly in promoter regions and show a high motif score and enrichment in both cell types (**Figure 4B**). We provide an illustrative example of cell type-specific TF binding sites for *TRPM3* (Transient Receptor Potential Cation Channel, Subfamily M, Member 3), where two CTCF sites are predicted only in neuronal samples (**Figure 4C**).

**Figure 4.**
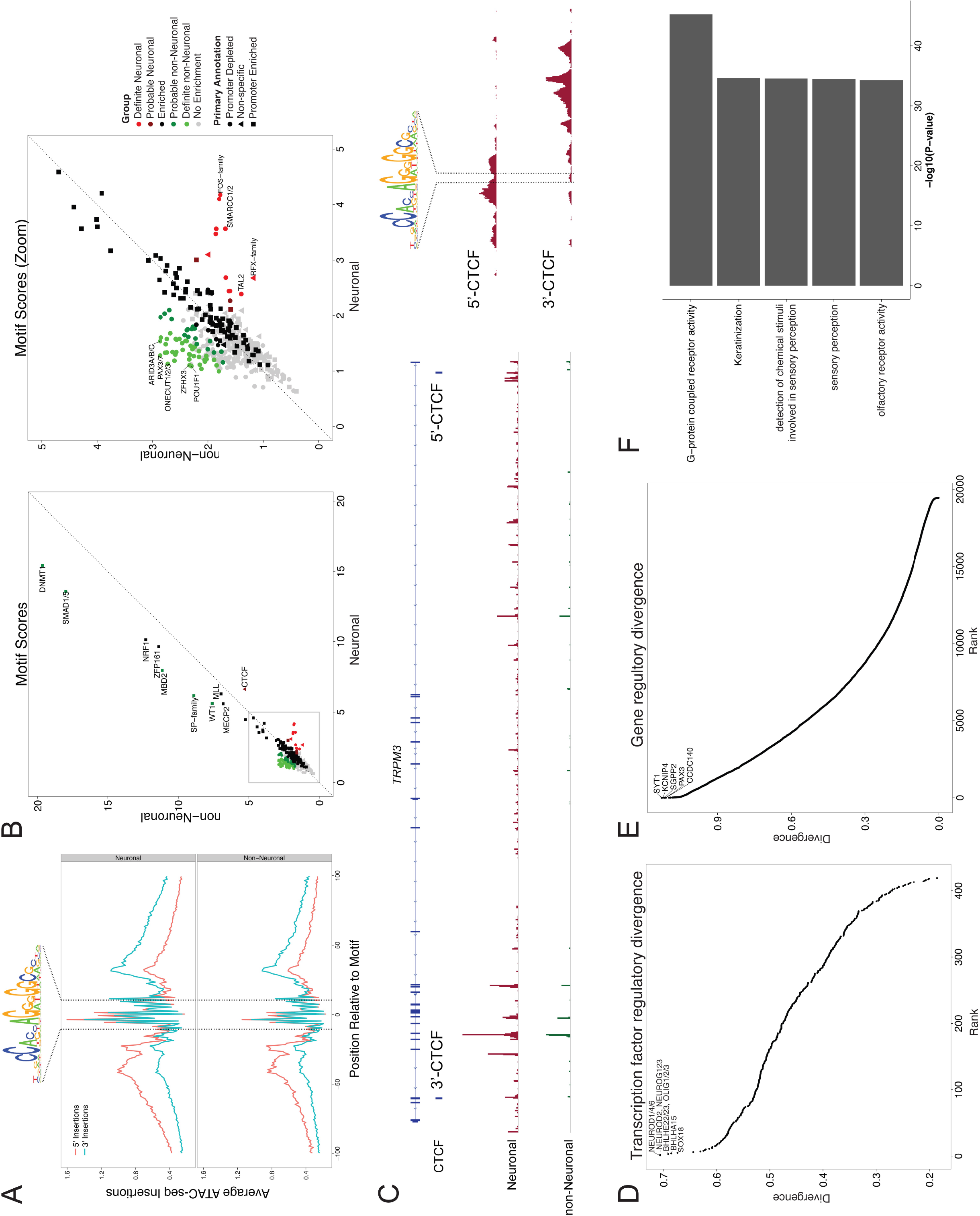
Transcription factor and gene regulome in neuronal and non-neuronal cells. **(A)** The average transposase insertion probability at all predicted binding CTCF binding sites within neuronal and non-neuronal OCRs. **(B)** Motif scores in neuronal and non-neuronal cells. Based on the ratio of the motif scores between the two samples, the cell-type specificity of the motifs is categorized as probable (≥1.25) and definite (≥1.5). Motifs with an enrichment of matches within motifs compared to the genomic background of less than 1.25 were categorized with “no enrichment”. Based on the relative occurrence of motifs in promoter peaks compared to all peaks, the motifs were categorized as promoter depleted (≤1/1.25) or enriched (≥1.25). We highlight the most enriched motifs for all TFs (Left panel). A zoomed version (Gray box) is illustrated on the right panel. Here, the most cell-type specific TFs are highlighted, listing only the most significant TF of each TF-family. **(C)** Two CTCF predicted sites near the 3’ and 5’ ends of the TRPM3 gene were detected only in neuronal cells. The exact location of the CTCF binding motif for the 5’-CTCF and 3’-CTCF binding sites is illustrated on the right. **(D)** The regulatory divergence of TF targets between the neuronal and non-neuronal samples. **(E)** The regulatory divergence of genes between the neuronal and non-neuronal samples. **(F)** Gene Ontology terms enriched in the top 1000 most divergently regulated genes.

Based on their relative proximity to TF binding sites identified by ATACseq, we next examined the likelihood that the TFs were regulating the same or different genes in the neuronal and non-neuronal samples. We did this by calculating the regulatory divergence (Qu et al., 2015) between these two cell types (**Supplemental Experimental Procedures**). The two most divergent motifs jointly represented the neurogenin and neuroD TF families (**Figure 4D**). These TFs are, therefore, predicted to regulate a different set of genes in neuronal and non-neuronal cells. At the other end of the spectrum, the least divergent TFs consisted of many general TFs, including NFYB and NFYA (**Table S5**). As an alternative approach, we applied the concept of regulatory divergence to identify genes that showed either a similar or a dissimilar pattern of TF regulation in the two cell types. Several genes with marked regulatory differences between neurons and non-neurons were identified (**Figure 4E**). To ensure that these estimated gene regulatory divergences were not erroneously driven by multiple overlapping, similar motifs, we created a non-redundant set of motifs, each with less than 25% overlap to any other motif, and re-ran the analysis using these. This yielded highly similar results (data not shown). The most divergently regulated gene was *SYT1*, which encodes Synaptotagmin 1, a neuronally expressed protein known to mediate synaptic vesicle exocytosis (Lee et al., 2010). Pathway analysis of the top 1000 most divergently regulated genes showed significant enrichment for biological processes related to G-protein coupled receptor activity (**Figure 4F**).

### Enrichment of SCZ risk loci for OCRs and TF binding sites

We sought to determine if the genomic regions corresponding to identified OCRs contain common risk variants detected through genome wide association studies (GWAS). As an initial analysis, we tested if the identified OCRs were enriched in risk loci for SCZ (PGC-SCZ, 2014), Alzheimer’s disease (AD) (Lambert et al., 2013) and non-neuropsychiatric diseases, including inflammatory bowel disease (Liu et al., 2015), rheumatoid arthritis (Okada et al., 2014), coronary artery disease (Consortium, 2015) and lipid traits (Global Lipids Genetics, 2013). For the analysis, described below, the level of enrichment of each functional annotation with GWAS traits was estimated using an empirical Bayes approach (Pickrell, 2014). We found enrichment for SCZ and neuronal [log2 enrichment (95% Confidence Interval or 95CI) × 2.11 (1.21 – 2.73)] and non-neuronal [log2 enrichment (95CI) = 1.57 (0.01 – 2.38] OCRs (**Figure S5A**). This is consistent with the tissue specific enrichment of OCRs with relevant diseases (Maurano et al., 2012; Roussos et al., 2014). In addition, compared to non-neuronal OCRs, we found a stronger enrichment in neuronal OCRs, thereby providing additional support for neurons as the functional unit affected by SCZ susceptibility loci.

Given the significant enrichment of OCRs with SCZ loci, we next tested whether OCRs within specific genic annotations are more enriched in SCZ. The most enriched annotations were neuronal introns [log2 enrichment (95CI) = 2.95 (2.10 – 3.55)] and non-neuronal promoters [log2 enrichment (95CI) = 3.33 (2.38 – 3.98)] (**Figure S5B**). These two annotations were used to build a combined model. We then used this combined model in a statistical approach to reweigh the SCZ GWAS and identify the functional variant underlying each disease-associated locus (**Supplementary Experimental Procedures)**. This yielded 29 SNPs that localize within 20 out of the 108 GWAS SCZ loci as the most likely candidates to be the causal polymorphisms in each region (**Table S6 OCR model**). For 19 out of the 20 GWAS SCZ loci, the functional SNP is not the GWAS index SNP. We found a substantial increase of ~10 folds for the likelihood (estimated based on the fitted empirical prior probability) of a functional SNP (average prior = 0.38%) to be the causal polymorphism in this region compared to the index GWAS SNP (average prior = 0.036%). We note, however, that the likelihood of the functional SNPs in each region is low (all priors < 1%). For 13 out of the 20 SCZ risk loci, we also identified an effect of the putative functional SNP on gene expression using expression quantitative trait analysis from the CommonMind Consortium (Fromer et al., 2016).

We then examined the enrichment of predicted TF sites within SCZ risk loci and identified 15 TFs (single TF model) derived mostly from neuronal (11 TFs) compared to non-neuronal (4 TFs) cells that were enriched with SCZ genetic variants (**Figure 5A**). These annotations were combined in a joint model, using cross-validation to overcome overfitting in the model (**Supplementary Experimental Procedures)**. Our best-fitting model included 4 TFs (combined TF model) ZSCAN10, NANOG/NANOGP1, CEBPZ and ZNF354C, all of which were specific to neuronal cells (**Figure 5A**). The single and combined TF models were used to reweigh the SCZ GWAS and identified 7 SNPs that localize within 6 out of the 108 GWAS SCZ loci as the candidates most likely to be the causal polymorphisms in each region (**Table S6 TF model**). In this analysis, none of the functional SNPs were the GWAS index SNP. We found a substantial increase of ~100 folds for likelihood (estimated based on the fitted empirical prior probability) of a functional SNP (average prior = 1.91%) to be the causal polymorphism in this region compared to the index GWAS SNP (average prior = 0.019%). For 3 out of the 6 SCZ risk loci, we also identified an effect of the putative functional SNP on gene expression using expression quantitative trait analysis from the CommonMind Consortium (Fromer et al., 2016).

**Figure 5.**
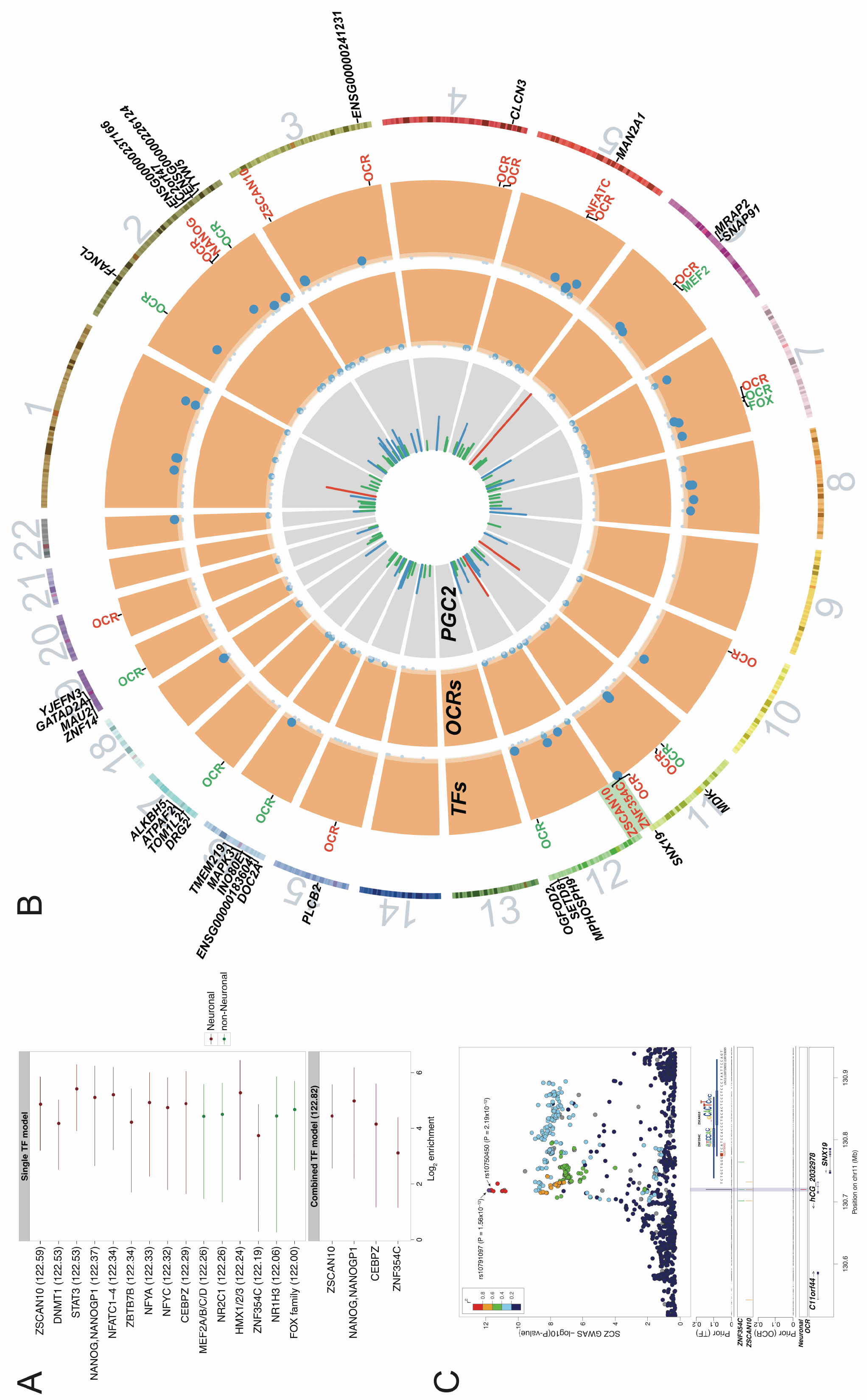
Integration of OCRs and TF binding sites with SCZ risk variants. **(A)** Enrichment of predicted TF sites within SCZ risk loci from neuronal and non-neuronal cells using a single TF model (top panel) and a joint model (bottom panel). The maximum-likelihood estimates and 95% confidence intervals of the enrichment parameter for each TF is illustrated. Annotations are ranked based on the improvement of the likelihood of the model (at the top are those that improved the likelihood the most). The number in parenthesis is the likelihood of the model for each TF (single model) or the combined model. **(B)** Circos plot showing the distribution of the fitted empirical prior probability that each SNP is the causal one in the 108 SCZ risk loci based on the OCR or TF (single and combined) models. The different layers of the circus plot show (from inside to outside) are: PGC2 SCZ risk loci (layer 1): the 108 loci SCZ loci and the level of significance for the index SNP of each locus is provided. Green lines illustrate loci with −log10 P value ≤ 10, blue lines show loci −log10 P value between 10 and 15 and red lines show loci with −log10 P value > 15. Fitted empirical prior probabilities for the OCR (layer 2) and TF (layer 3) models: SNPs with prior ≤ 0.002 are in light orange background and light blue dots; SNPs with 0.002 < prior ≤ 0.01 are in orange background and blue dots; SNPs with prior > 0.01 are in dark orange background and dark blue dots. Only the TF model detects functional SNPs with prior > 0.01. In layer 4, we illustrate whether a PGC2 locus has a functional SNP with a posterior probability of association (PPA) > 0.1 that overlaps an OCR or TF binding site. Red and green fonts are OCRs and TFs derived from the neuronal and non-neuronal cells, respectively. The green box indicates the locus detected by the combined model further illustrated in panel **(C)** Layer 5 shows genes with an eQTL effect for the functional SNPs. **(C)** Regional plot surrounding the *SNX19* locus. The top panel shows a plot of the *r^2^* and P values for association with SCZ, including the index (rs10791097) and the functional (rs10750450) SNP. In the middle panel (Prior TF) is the fitted empirical prior probability based on the TF combined model and the positions of the TFs for this region. The overlap of the SNP with the binding sites of ZSCAN10 and ZNF354C are illustrated. In the lower panel (Prior OCR) is the fitted empirical prior probability and position of the OCR model. Note the difference in the fitted empirical prior probability among the TF and OCR model for the functional SNP (in gray box).

**Figure 5B** shows the distribution of the prior probability of functional SNPs across the 108 GWAS SCZ loci. Compared to the neuronal introns and non-neuronal promoters combined model, the TF models (single and combined) include multiple functional SNPs with high likelihood (priors > 1%) in SCZ loci (**Figure 5B**). A subset of variants has additional support for a functional role, affecting gene expression abundance of specific transcripts. **Figure 5C** shows an illustrative example for a locus near the *SNX19* gene, where the combined TF model identified rs10750450 as the most likely causal variant in this region. This SNP has a P value of 2.19 × 10^−12^, which is close to the level of significance of the index SNP of that locus (rs10791097; P = 1.56 × 10^−12^). However, this SNP falls in the proximity of 2 TF binding sites (ZSCAN10 and ZNF354C), leading the model to assign a prior (14.58%) that is almost three orders of magnitude higher than the prior of the index SNP (0.02%). Overall, using the cell type-specific OCR and TF data, we assign a putative functional role for 20 and 6 SCZ risk loci (25 loci in total), respectively. Furthermore, we provide evidence that, compared to OCR, TF analysis defines a smaller number of risk loci with but with higher priors.

Finally, a recent study showed that synonymous, rare de novo mutations in SCZ, but not Autism spectrum disorders (ASD), are enriched for frontal cortex-derived DNase I hypersensitivity sites (Takata et al., 2016). We examined whether the ATAC-seq peaks are enriched in SCZ and ASD de novo synonymous mutations. Sixteen out of 228 de novo SCZ variants overlapped with an ATAC-seq peak in either neuron or non-neuronal cells compared to 4 out of 154 variants found in controls, which represents a significant enrichment (OR (95CI) = 2.83 (0.93 – 8.64); one-sided Fisher’s exact test P = 0.044) (**Table S7**). No significant enrichment was seen for the peaks specific to the neuronal (onesided Fisher’s exact test P = 0.54) and non-neuronal cells (one-sided Fisher’s exact test P = 0.21), potentially due to a lack of power. Similar to previous findings (Takata et al., 2016), we found no enrichment for ASD variants (one-sided Fisher’s exact test P = 0.41). Our results are consistent with the original study (Takata et al., 2016) and provide additional evidence that synonymous de novo mutations might play a significant role in SCZ by altering OCRs.

## DISCUSSION

Recent genetic studies have implicated numerous common risk variants in SCZ (PGC-SCZ, 2014). One of the next challenges is to further understand the biological mechanisms of the large number, and diversity, of genes that are associated with SCZ. To that end, we need to generate additional data capturing putative molecular processes that are relevant to the development of the disease. The majority of SCZ risk variants is found within non-coding regions of the genome and is predicted to disrupt the function of CREs. We further explored the SCZ genetic architecture by leveraging, for the first time to our knowledge, cell type-specific OCRs mapped in human brain tissue. While, SCZ-associated abnormalities have been demonstrated in neuronal and non-neuronal cells (Insel, 2010; Roussos and Haroutunian, 2014), here we demonstrate that SCZ risk variants show a higher enrichment in neuronal OCRs. This is consistent with genetic findings implicating genes that participate in neuronal function and synaptic transmission in the etiology of SCZ (Fromer et al., 2014; PGC-SCZ, 2014; Purcell et al., 2014).

To further refine the OCRs, we performed TF digital footprinting analysis, which provides higher resolution of the functional genomic regions (from ~1kb average OCR size to ~10bp for TF binding site) and assigns a potential functional role based on known TF. Using the TF compared to the OCR data, we found a substantial increase of ~5 folds for likelihood of a functional SNP to be the causal polymorphism in certain SCZ risk regions. In addition, we identified a subset of TFs that are highly enriched in SCZ loci, including ZSCAN10, NANOG/NANOGP1, CEBPZ and ZNF354C, all of which were specific to neuronal cells. While little is known about the function of CEBPZ and ZNF354C, both NANOG and ZSCAN10 have been shown to play a role in the maintenance of pluripotency in embryonic stem cells (ESCs). NANOG is a downstream target of ZSCAN10 transcriptional activity (Wang et al., 2007) and NANOG expression is thought to be restricted to ESCs, becoming progressively down-regulated during differentiation and embryonic development (Mitsui et al., 2003; Thomson et al., 2011). This raises the question as to why we detect TF footprints for developmental genes in neurons of the adult brain? While it is possible that NANOG and ZSCAN10 have, heretofore unknown, roles in postmitotic neurons, recent evidence suggests that pro-neural TF footprints can be maintained over time to ensure proper neuronal and glial differentiation. As such, we may be detecting the impression left by a protein on the structure of DNA, several decades after the fact (Ziller et al., 2015). To that end, there is evidence supporting a role for ZSCAN10 in maintaining a multipotent progenitor cell population in mid-gestation embryos and adult organs (Kraus et al., 2014).

The SCZ risk regions are frequently large and often contain multiple implicated SNPs due to local linkage disequilibrium patterns. In order to be able to understand these associations mechanistically, it is important to develop strategies for honing in on regions and SNPs more likely to have functional effects. We applied our TF footprinting models to reweigh the SCZ GWAS and identified the variants that have a putative functional role for 6 out of the 108 GWAS SCZ risk loci, none of which is the index SNP identified by GWAS. One such example is a locus adjacent to the *SNX19* gene where the index SNP for this locus was identified as rs10791097. Our combined TF model identified a different SNP, rs10750450, as the most likely causal variant in this region, due to its proximity to binding sites for ZSCAN10 and ZNF354C in neuronal cells. This SNP is an eQTL for the *SNX19* transcript in the human brain (based on CMC data), but also in multiple other tissues based on the GTEx data (http://www.gtexportal.org/). Interestingly, SNX19 was recently identified as one (out of two genes) whose expression level was associated with SCZ (Zhu et al., 2016). Here we add to the growing evidence that upregulation of SNX19 increases the risk of SCZ by localizing the functional regulatory region in a specific cell type.

Although we observed an increased frequency of accessible elements for both cell-types in proximity to transcription start sites (TSSs), distal regulatory regions appear to play a more critical role in neurons when compared to non-neurons. This finding compliments a previous study suggesting that, compared to non-neurons, neuron specific methylated regions tend to be located distally from TSSs and are enriched within predicted enhancer elements (Kozlenkov et al., 2014). Furthermore, while both neuronal and non-neuronal OCRs are evolutionary conserved, those of neurons appear to be under stronger functional constraint than other brain cells. We also observe cell-type differences in the regulation of gene expression between the two cell types. Promoter-depleted TFs showed the most marked differences between cells. Based on our analysis, the TFs with higher binding affinity for neurons were among the FOS/JUN (Raivich and Behrens, 2006) and bHLH families (Ross et al., 2003) whereas PAX3/7 (Monsoro-Burq, 2015) and the homeodomain/sox (Reiprich and Wegner, 2015) TF families display non-neuronal preference.

XCI involves the heterochromatic silencing of one copy of the X chromosome in the cells of female mammals. A subset of X-linked genes escape XCI and, for some genes, this occurs in a tissue and cell type specific manner (Deng et al., 2014). X□linked forms of intellectual disability are 3.5 times more common than autosomal forms. ID and autism are more prevalent in males. Females are thought to be protected from disease due to having a mixture of cells expressing different sets of X-linked genes through XCI skewing or escape (Deng et al., 2014). The proportion of X□linked genes expressed in the brain is significantly higher than in other somatic tissues and approximately 100 human X□linked genes associated with intellectual disability have been identified (Deng et al., 2014). Interestingly, the effects of these mutations are more variable in females due to genes that escape XCI. Using our OCR data, we implicate cell-type specific OCRs in XCI, including regions that have been previously associated with XLID.

Our study is not without limitations, including a relatively small sample size, and these results warrant further validation in future studies. As is the case with most postmortem studies, the current observations could be partially attributed to technical and clinical covariates, including postmortem interval and agonal state. In addition, we chose to focus on a discreet region of the brain and, due to the limitations of working with frozen postmortem tissue, and the loss of cytoplasmic markers upon thawing, restricted our analysis to the study of two broad populations of cells – neurons and nonneurons. The study of different regions of the brain coupled with the application of additional cell-type specific nuclear markers e.g. (Kozlenkov et al., 2015) would broaden the scope of our approach to reach a more thorough understanding of the means by which CREs influence brain function and disease.

We have generated the first, cell-type-specific, open chromatin maps of human postmortem brain. This has allowed us to identify specific patterns of gene regulation in neuronal and non-neuronal cells and to assign functional roles to non-coding SCZ risk variants.

## EXPERIMENTAL PROCEDURES

See also Supplemental Experimental Procedures.

### Brain Samples and ATAC-seq libraries

Brain tissue specimens were obtained from the NIH Brain and Tissue Repository at Icahn School of Medicine at Mount Sinai and JJ Peters VA Medical Center. We processed 50mg of tissue from the frontopolar prefrontal cortex of 8 controls (**Table 1**). DAPI positive neuronal (NeuN+) and nonneuronal (NeuN−) nuclei were sorted using a FACSAria flow cytometer (BD Biosciences). ATAC-seq reactions were performed using an established protocol (Buenrostro et al., 2013) with minor modifications. Libraries were amplified for a total of 9-14 cycles and were quantified by Qubit HS DNA kit (Life technologies) and by quantitiative PCR (KAPA Biosystems Cat#KK4873) prior to sequencing. Libraries were sequenced on Hi-Seq2500 (Illumina) obtaining 2×50 paired-end reads.

### Data processing and differential analysis

Reads were mapped to the gender appropriate reference genome hg19 using STAR aligner (Dobin et al., 2013) version 2.5.0. For each sample, this produced a coordinate-sorted BAM file of mapped paired-end reads. We excluded reads that: (1) mapped to more than one locus; (2) were duplicated; and/or (3) mapped to the mitochondrial genome. Peaks were called using MACS2 v2.1 (Zhang et al., 2008). Peak sets across cell types were consolidated into two single lists by union operations of peaks that were present in at least two libraries. The final consensus set of 115,021 peaks was generated by combining the neuronal and non-neuronal peaks. We used the featureCounts function from the Rsubread package (Liao et al., 2014) to generate a sample-by-peak read count matrix (16 samples by 151,021 peak regions). We used the edgeR package (Robinson et al., 2010) to model the normalized read counts using negative binomial (NB) distributions including cell type, gender, age of death and PMI as covariates. A quasi-likelihood (QL) F-test was conducted for each OCR using the glmQLFTest function (Lund et al., 2012). p-values were then adjusted for multiple hypothesis testing using false discovery rate (FDR) below, or at, 1%.

### Transcription factor analysis

All transcription factor motifs representing human transcription factors were downloaded from the meta-database CIS-BP 1.02 (Weirauch et al., 2014). PIQ (Sherwood et al., 2014) was then used to predict transcription factor binding sites from the genome sequence. Using the PIQ genome-wide predicted TF binding sites, we calculated the relative occurrence of motifs within the peaks ("peak enrichments") as: motifs in ATAC-peaks per bp divided by motifs in the genome per genome size. For each motif, we retained only binding sites that were within the ATAC-seq peaks and passed the default purity cut-off (70%).

### Integrating functional annotations with GWAS

To integrate functional annotations and GWAS results, we used the fGWAS software (Pickrell, 2014), which implements an empirical Bayesian framework to identify genomic annotations that are enriched, or depleted, for loci influencing a trait. This algorithm estimates the prior probability that a given block contains an association and the conditional prior probability that a given SNP in the block is causal, based on the presence of functional annotations. We performed two different models that considered OCRs (OCR model) or TF binding sites (TF model). Functional SNPs were further explored using expression quantitative trait loci (eQTLs) from prefrontal cortex (Fromer et al., 2016).

## AUTHOR CONTRIBUTIONS

J.F.F. and P.R. conceived the project and designed all experiments. V.H. performed the tissue dissections. J.F.F. and C.B. performed the FANS. J.F.F. prepared the sequencing libraries. C.G., M.E.H., K.X., J.T.D., M.M., J.K.P. and P.R. advised on analysis approaches and analyzed data. J.F.F., C.G., M.E.H. and P.R. wrote the manuscript.

## ACKNOWLEDGMENTS

This work was supported by the National Institutes of Health (R01AG050986 Roussos), Brain Behavior Research Foundation (20540 Roussos), Alzheimer’s Association (NIRG-340998 Roussos) and the Veterans Affairs (Merit grant BX002395 Roussos).

## References

Andersson, R., Gebhard, C., Miguel-Escalada, I., Hoof, I., Bornholdt, J., Boyd, M., Chen, Y., Zhao, X., Schmidl, C., Suzuki, T., et al. (2014). An atlas of active enhancers across human cell types and tissues. Nature 507, 455–461.

Baldwin, C.T., Hoth, C.F., Macina, R.A., and Milunsky, A. (1995). Mutations in PAX3 that cause Waardenburg syndrome type I: ten new mutations and review of the literature. American journal of medical genetics 58, 115–122.

Bernstein, B.E., Birney, E., Dunham, I., Green, E.D., Gunter, C., and Snyder, M. (2012). An integrated encyclopedia of DNA elements in the human genome. Nature 489, 57–74.

Buenrostro, J.D., Giresi, P.G., Zaba, L.C., Chang, H.Y., and Greenleaf, W.J. (2013). Transposition of native chromatin for fast and sensitive epigenomic profiling of open chromatin, DNA-binding proteins and nucleosome position. Nature methods 10, 1213–1218.

Consortium, C.D. (2015). A comprehensive 1000 Genomes-based genome-wide association metaanalysis of coronary artery disease. Nat Genet 47, 1121–1130.

Cooper, G.M., Stone, E.A., Asimenos, G., Program, N.C.S., Green, E.D., Batzoglou, S., and Sidow, A. (2005). Distribution and intensity of constraint in mammalian genomic sequence. Genome Res 15, 901–913.

Deng, X., Berletch, J.B., Nguyen, D.K., and Disteche, C.M. (2014). X chromosome regulation: diverse patterns in development, tissues and disease. Nature reviews Genetics 15, 367–378.

Dobin, A., Davis, C.A., Schlesinger, F., Drenkow, J., Zaleski, C., Jha, S., Batut, P., Chaisson, M., and Gingeras, T.R. (2013). STAR: ultrafast universal RNA-seq aligner. Bioinformatics 29, 15–21.

Fromer, M., Pocklington, A.J., Kavanagh, D.H., Williams, H.J., Dwyer, S., Gormley, P., Georgieva, L., Rees, E., Palta, P., Ruderfer, D.M., et al. (2014). De novo mutations in schizophrenia implicate synaptic networks. Nature 506, 179–184.

Fromer, M., Roussos, P., Sieberts, S.K., Johnson, J.S., Kavanagh, D.H., Perumal, T.M., Ruderfer, D.M., Oh, E.C., Topol, A., Shah, HR., et al. (2016). Gene Expression Elucidates Functional Impact of Polygenic Risk for Schizophrenia. bioRxiv.

Fullard, J.F., Halene, T.B., Giambartolomei, C., Haroutunian, V., Akbarian, S., and Roussos, P. Understanding the genetic liability to schizophrenia through the neuroepigenome. Schizophrenia research.

Gilbert, S.J., Spengler, S., Simons, J.S., Steele, J.D., Lawrie, S.M., Frith, C.D., and Burgess, P.W. (2006). Functional Specialization within Rostral Prefrontal Cortex (Area 10): A Meta-analysis. Journal of Cognitive Neuroscience 18, 932–948.

Global Lipids Genetics, C. (2013). Discovery and refinement of loci associated with lipid levels. Nat Genet 45, 1274–1283.

Hacisuleyman, E., Goff, L.A., Trapnell, C., Williams, A., Henao-Mejia, J., Sun, L., McClanahan, P.,Hendrickson, D.G., Sauvageau, M., Kelley, DR., et al. (2014). Topological organization of multichromosomal regions by the long intergenic noncoding RNA FIRRE. Nature structural & molecular biology 21, 198–206.

Insel, T.R. (2010). Rethinking schizophrenia. Nature 468, 187–193.

Kozlenkov, A., Roussos, P., Timashpolsky, A., Barbu, M., Rudchenko, S., Bibikova, M., Klotzle, B., Byne, W., Lyddon, R., Di Narzo, A.F., et al. (2014). Differences in DNA methylation between human neuronal and glial cells are concentrated in enhancers and non-CpG sites. Nucleic acids research 42, 109–127.

Kozlenkov, A., Wang, M., Roussos, P., Rudchenko, S., Barbu, M., Bibikova, M., Klotzle, B., Dwork, A.J., Zhang, B., Hurd, Y.L., et al. (2015). Substantial DNA methylation differences between two major neuronal subtypes in human brain. Nucleic acids research.

Kraus, P., V. S. Yu, H.B., Xing, X., Lim, S.L., Adler, T., Pimentel, J.A.A., Becker, L., Bohla, A., Garrett, L., et al. (2014). Pleiotropic Functions for Transcription Factor Zscan10. PLoS ONE 9, e104568.

Lambert, J.C., Ibrahim-Verbaas, C.A., Harold, D., Naj, A.C., Sims, R., Bellenguez, C., DeStafano, A.L., Bis, J.C., Beecham, G.W., Grenier-Boley, B., et al. (2013). Meta-analysis of 74,046 individuals identifies 11 new susceptibility loci for Alzheimer’s disease. Nat Genet 45, 1452–1458.

Lee, H-K., Yang, Y., Su, Z., Hyeon, C., Lee, T.-S., Lee, H.-W., Kweon, D.-H., Shin, Y.-K., and Yoon, T.-Y. (2010). Dynamic Ca2+-dependent stimulation of vesicle fusion by membrane-anchored synaptotagmin 1. Science 328, 760–763.

Lee, J.T., and Bartolomei, M.S. (2013). X-inactivation, imprinting, and long noncoding RNAs in health and disease. Cell 152, 1308–1323.

Liao, Y., Smyth, G.K., and Shi, W. (2014). featureCounts: an efficient general purpose program for assigning sequence reads to genomic features. Bioinformatics 30, 923–930.

Liu, J.Z., van Sommeren, S., Huang, H., Ng, S.C., Alberts, R., Takahashi, A., Ripke, S., Lee, J.C., Jostins, L., Shah, T., et al. (2015). Association analyses identify 38 susceptibility loci for inflammatory bowel disease and highlight shared genetic risk across populations. Nat Genet 47, 979–986.

Lund, S.P., Nettleton, D., McCarthy, D.J., and Smyth, G.K. (2012). Detecting differential expression in RNA-sequence data using quasi-likelihood with shrunken dispersion estimates. Statistical applications in genetics and molecular biology 11.

Maurano, M.T., Humbert, R., Rynes, E., Thurman, R.E., Haugen, E., Wang, H., Reynolds, A.P., Sandstrom, R., Qu, H., Brody, J., et al. (2012). Systematic localization of common disease-associated variation in regulatory DNA. Science 337, 1190–1195.

Mitsui, K., Tokuzawa, Y., Itoh, H., Segawa, K., Murakami, M., Takahashi, K., Maruyama, M., Maeda, M., and Yamanaka, S. (2003). The Homeoprotein Nanog Is Required for Maintenance of Pluripotency in Mouse Epiblast and ES Cells. Cell 113, 631–642.

Mo, A., Mukamel, E.A., Davis, F.P., Luo, C., Henry, G.L., Picard, S., Urich, M.A., Nery, J.R., Sejnowski, T.J., Lister, R., et al. (2015). Epigenomic Signatures of Neuronal Diversity in the Mammalian Brain. Neuron 86, 1369–1384.

Monsoro-Burq, A.H. (2015). PAX transcription factors in neural crest development. Seminars in cell & developmental biology 44, 87–96.

Neph, S., Vierstra, J., Stergachis, A.B., Reynolds, A.P., Haugen, E., Vernot, B., Thurman, R.E., John, S., Sandstrom, R., and Johnson, A.K. (2012). An expansive human regulatory lexicon encoded in transcription factor footprints. Nature 489, 83–90.

Okada, Y., Wu, D., Trynka, G., Raj, T., Terao, C., Ikari, K., Kochi, Y., Ohmura, K., Suzuki, A., Yoshida, S., et al. (2014). Genetics of rheumatoid arthritis contributes to biology and drug discovery. Nature 506, 376–381.

PGC-SCZ (2014). Biological insights from 108 schizophrenia-associated genetic loci. Nature 511, 421–427.

Pickrell, J.K. (2014). Joint analysis of functional genomic data and genome-wide association studies of 18 human traits. American journal of human genetics 94, 559–573.

Purcell, S.M., Moran, J.L., Fromer, M., Ruderfer, D., Solovieff, N., Roussos, P., O’Dushlaine, C., Chambert, K., Bergen, S.E., Kahler, A., et al. (2014). A polygenic burden of rare disruptive mutations in schizophrenia. Nature 506, 185–190.

Purohit, D.P., Perl, D.P., Haroutunian, V., Powchik, P., Davidson, M., and Davis, K.L. (1998). Alzheimer disease and related neurodegenerative diseases in elderly patients with schizophrenia: a postmortem neuropathologic study of 100 cases. Archives of general psychiatry 55, 205–211.

Qu, K., Zaba, Lisa C., Giresi, Paul G., Li, R., Longmire, M., Kim, Youn H., Greenleaf, William J., and Chang, Howard Y. Individuality and Variation of Personal Regulomes in Primary Human T Cells. Cell Systems 1, 51–61.

Qu, K., Zaba, L.C., Giresi, P.G., Li, R., Longmire, M., Kim, Y.H., Greenleaf, W.J., and Chang, H.Y. (2015). Individuality and variation of personal regulomes in primary human T cells. Cell Systems 1, 51–61.

Raivich, G., and Behrens, A. (2006). Role of the AP-1 transcription factor c-Jun in developing, adult and injured brain. Progress in neurobiology 78, 347–363.

Reiprich, S., and Wegner, M. (2015). From CNS stem cells to neurons and glia: Sox for everyone. Cell and tissue research 359, 111–124.

Roadmap Epigenomics, C., Kundaje, A., Meuleman, W., Ernst, J., Bilenky, M., Yen, A., Heravi-Moussavi, A., Kheradpour, P., Zhang, Z., Wang, J., et al. (2015). Integrative analysis of 111 reference human epigenomes. Nature 518, 317–330.

Robinson, M.D., McCarthy, D.J., and Smyth, G.K. (2010). edgeR: a Bioconductor package for differential expression analysis of digital gene expression data. Bioinformatics 26, 139–140.

Ross, S.E., Greenberg, M.E., and Stiles, C.D. (2003). Basic Helix-Loop-Helix Factors in Cortical Development. Neuron 39, 13–25.

Roussos, P., and Haroutunian, V. (2014). Schizophrenia: susceptibility genes and oligodendroglial and myelin related abnormalities. Frontiers in Cellular Neuroscience 8.

Roussos, P., Mitchell, A.C., Voloudakis, G., Fullard, J.F., Pothula, V.M., Tsang, J., Stahl, E.A., Georgakopoulos, A., Ruderfer, D.M., Charney, A., et al. (2014). A role for noncoding variation in schizophrenia. Cell Rep 9, 1417–1429.

Seo, H.-C., Sætre, B.O., Håvik, B., Ellingsen, S., and Fjose, A. (1998). The zebrafish Pax3 and Pax7 homologues are highly conserved, encode multiple isoforms and show dynamic segment-like expression in the developing brain. Mechanisms of development 70, 49–63.

Sherwood, R.I., Hashimoto, T., O’Donnell, C.W., Lewis, S., Barkal, A.A., van Hoff, J.P., Karun, V., Jaakkola, T., and Gifford, D.K. (2014). Discovery of directional and nondirectional pioneer transcription factors by modeling DNase profile magnitude and shape. Nature biotechnology 32, 171-178.

Takata, A., Ionita-Laza, I., Gogos, Joseph A., Xu, B., and Karayiorgou, M. (2016). De Novo Synonymous Mutations in Regulatory Elements Contribute to the Genetic Etiology of Autism and Schizophrenia. Neuron 89, 940–947.

Takizawa, R., Kasai, K., Kawakubo, Y., Marumo, K., Kawasaki, S., Yamasue, H., and Fukuda, M. Reduced frontopolar activation during verbal fluency task in schizophrenia: A multi-channel near-infrared spectroscopy study. Schizophrenia research 99, 250–262.

Thomson, M., Liu, Siyuan J., Zou, L.-N., Smith, Z., Meissner, A., and Ramanathan, S. (2011). Pluripotency Factors in Embryonic Stem Cells Regulate Differentiation into Germ Layers. Cell 145, 875–889.

Thurman, R.E., Rynes, E., Humbert, R., Vierstra, J., Maurano, M.T., Haugen, E., Sheffield, N.C., Stergachis, A.B., Wang, H., and Vernot, B. (2012). The accessible chromatin landscape of the human genome. Nature 489, 75–82.

Trynka, G., Sandor, C., Han, B., Xu, H., Stranger, B.E., Liu, X.S., and Raychaudhuri, S. (2013). Chromatin marks identify critical cell types for fine mapping complex trait variants. Nat Genet 45, 124-130.

Wang, Z.-X., Kueh, J.L.L., Teh, C.H.-L., Rossbach, M., Lim, L., Li, P., Wong, K.-Y., Lufkin, T., Robson, P., and Stanton, L.W. (2007). Zfp206 Is a Transcription Factor That Controls Pluripotency of Embryonic Stem Cells. STEM CELLS 25, 2173–2182.

Weirauch, M.T., Yang, A., Albu, M., Cote, A.G., Montenegro-Montero, A., Drewe, P., Najafabadi, H.S., Lambert, S.A., Mann, I., and Cook, K. (2014). Determination and inference of eukaryotic transcription factor sequence specificity. Cell 158, 1431–1443.

Yang, F., Deng, X., Ma, W., Berletch, J.B., Rabaia, N., Wei, G., Moore, J.M., Filippova, G.N., Xu, J., Liu, Y., et al. (2015). The lncRNA FIRRE anchors the inactive X chromosome to the nucleolus by binding CTCF and maintains H3K27me3 methylation. Genome Biol 16, 52.

Zeisel, A., Munoz-Manchado, A.B., Codeluppi, S., Lonnerberg, P., La Manno, G., Jureus, A., Marques, S., Munguba, H., He, L., Betsholtz, C., et al. (2015). Brain structure. Cell types in the mouse cortex and hippocampus revealed by single-cell RNA-seq. Science 347, 1138–1142.

Zhang, Y., Chen, K., Sloan, S.A., Bennett, M.L., Scholze, A.R., O’Keeffe, S., Phatnani, H.P., Guarnieri, P., Caneda, C., Ruderisch, N., et al. (2014). An RNA-Sequencing Transcriptome and Splicing Database of Glia, Neurons, and Vascular Cells of the Cerebral Cortex. The Journal of neuroscience: the official journal of the Society for Neuroscience 34, 11929–11947.

Zhang, Y., Liu, T., Meyer, C.A., Eeckhoute, J., Johnson, D.S., Bernstein, B.E., Nusbaum, C., Myers, R.M., Brown, M., Li, W., et al. (2008). Model-based analysis of ChIP-Seq (MACS). Genome Biol 9, R137.

Zhang, Y., Morales, A.C., Jiang, M., Zhu, Y., Hu, L., Urrutia, A.O., Kong, X., and Hurst, L.D. (2013). Genes that escape X-inactivation in humans have high intraspecific variability in expression, are associated with mental impairment but are not slow evolving.Molecular biology and evolution.

Zhu, J., Adli, M., Zou, J.Y., Verstappen, G., Coyne, M., Zhang, X., Durham, T., Miri, M., Deshpande, V., De Jager, P.L., et al. (2013). Genome-wide Chromatin State Transitions Associated with Developmental and Environmental Cues. Cell 152, 642–654.

Zhu, Z., Zhang, F., Hu, H., Bakshi, A., Robinson, M.R., Powell, J.E., Montgomery, G.W., Goddard, M.E., Wray, N.R., Visscher, P.M., et al. (2016). Integration of summary data from GWAS and eQTL studies predicts complex trait gene targets. Nat Genet 48, 481–487.

Ziller, M.J., Edri, R., Yaffe, Y., Donaghey, J., Pop, R., Mallard, W., Issner, R., Gifford, C.A., Goren, A., Xing, J., et al. (2015). Dissecting neural differentiation regulatory networks through epigenetic footprinting. Nature 518, 355–359.

